# Population genomic SNPs from epigenetic RADs: gaining genetic and epigenetic data from a single established next-generation sequencing approach

**DOI:** 10.1101/737940

**Authors:** M. Crotti, C.E. Adams, K.R. Elmer

## Abstract

1. Epigenetics is increasingly recognised as an important molecular mechanism underlying phenotypic variation. To study DNA methylation in ecological and evolutionary contexts, epiRADseq is a cost-effective next-generation sequencing technique based on reduced representation sequencing of genomic regions surrounding non-/methylated sites. EpiRADseq for genome-wide methylation abundance and ddRADseq for genome-wide SNP genotyping follow very similar library and sequencing protocols, but to date these two types of dataset have been handled separately. Here we test the performance of using epiRADseq data to generate SNPs for population genomic analyses.
2. We tested the robustness of using epiRADseq data for population genomics with two independent datasets: a newly generated single-end dataset for the European whitefish *Coregonus lavaretus*, and a re-analysis of publicly available, previously published paired-end data on corals. Using standard bioinformatic pipelines with a reference genome and without (i.e. *de novo* catalogue loci), we compared the number of SNPs retained, population genetic summary statistics, and population genetic structure between data drawn from ddRADseq and epiRADseq library preparations.
3. We find that SNPs drawn from epiRADseq are similar in number to those drawn from ddRADseq, with a 55-83% of SNPs being identified by both methods. Genotyping error rate was <5% in both approaches. For summary statistics such as heterozygosity and nucleotide diversity, there is a strong correlation between methods (Spearman’s rho > 0.88). Furthermore, identical patterns of population genetic structure were recovered using SNPs from epiRADseq and ddRADseq approaches.
4. We show that SNPs obtained from epiRADseq are highly similar to those from ddRADseq and are equivalent for estimating genetic diversity and population structure. This finding is particularly relevant to researchers interested in genetics and epigenetics on the same individuals because using a single epigenomic approach to generate two datasets greatly reduces the time and financial costs compared to using these techniques separately. It also efficiently enables correction of epigenetic estimates with population genetic data. Many studies will benefit from a combinatorial approach with genetic and epigenetic markers and this demonstrates a single, efficient method to do so.

## Introduction

The advent of Next Generation Sequencing (NGS) has facilitated a revolution in ecology and evolution by enabling the integration of the two fields to better elucidate molecular patterns and mechanisms (Ekblom & Galindo, 2011). Technologically, an advance of NGS is not just reduced, per base pair costs of sequencing but that genomic techniques can be applied to so-called ‘non-model’ species or those without reference genomes or other genomic resources (Ekblom & Galindo, 2011). Among the many NGS techniques recently developed, genotyping by sequencing approaches, such as restriction site associated DNA sequencing (RADseq), have stood out for their versatility, low cost, and the amount of data generated (Davey et al. 2013; Andrews et al. 2016; Rowe, Renaut, & Guggisberg, 2016). Briefly, one or more restriction enzymes are used to digest the genome and only fragments in a specified range are retained for sequencing, resulting in genotypes from a representative portion of the genome for a variable number of individuals (Andrews et al. 2016). Double digest RADseq, or ddRADseq (Peterson et al. 2012), is one of the many varieties of genotyping by sequencing methods available and is particularly powerful because it allows a high degree of customisation in terms of the number of loci obtained and coverage per individual, and can be modified for different sequencing platforms (Puritz et al. 2014; Recknagel et al. 2016). ddRADseq is now an established tool for genotyping with NGS, to investigate many topics in ecology and evolution including population genetics, genetic mapping, parentage inference, genomics of adaptation, and phylogenomics using single nucleotide polymorphisms (SNPs) (Davey & Blaxter, 2010; Andrews et al. 2016). SNPs focus on genetic mutations, but it is well recognised that other molecular processes in the genome such as gene regulation and methylation influence biodiversity.

The study of epigenetic processes, which cause change in gene expression without nucleotide mutation of the underlying genome sequence, is providing a new complexity in the genotype – phenotype map and in some cases a disconnect of genotype and phenotype (Feil & Fraga, 2012). The best understood epigenetic mechanism is DNA methylation, which involves the addition of a methyl group to cytosine, and in eukaryotes it occurs mainly in CpG dinucleotides (Metzger & Schulte, 2016). Relevant for ecology and evolution, the field of ecological epigenetics aims to understand how DNA methylation associates with patterns of population variation and influences phenotypic diversity, local adaptation, and plasticity in natural populations (Bossdorf, Richards, & Pigliucci, 2008; Hu & Barrett, 2017). Until recently, epigenetic research in wild populations was conducted mainly using methylation-sensitive AFLPs (MS-AFLP) (e.g. Foust et al. 2016; Herrera et al. 2016), since they are cost-effective, easily applied to non-model organisms, and not computationally demanding (Schrey et al. 2013). However, they have several shortcomings (see review by Schrey et al. (2013)), the greatest of which is that they screen anonymous loci that then cannot be genome referenced nor compared across studies. Recently, the field has been invigorated by new methods that take advantage of NGS technology. One example is bisulfite sequencing, which comes in a number of variations (whole genome, reduced representation, target sequencing of specific gene regions) and has been shown to provide high resolution of the methylation landscape within genomes (Metzger & Schulte, 2016). However, this technique is expensive, can result in excessive DNA degradation, and requires a related reference genome for the species of interest, something that is still lacking for most non-model organisms (Leontiou et al. 2015; Metzger & Schulte, 2016).

EpiRADseq is a recently developed, reduced representation approach (Schield et al. 2016) to study DNA methylation variation in individuals. It is based on the established ddRADseq protocol (Peterson et al. 2012) and involves the digestion of the genome using two restriction enzymes, with one enzyme being methylation-sensitive. Therefore, a methylated locus will not be cut by the methylation-sensitive enzyme, will not be enriched by PCR nor sequenced, and thus no sequencing read is obtained in the data. If a locus is unmethylated, it will be cut in the same way as ddRADseq and therefore enriched by PCR and sequenced. Therefore, the number of overall reads for a locus is proportional to the level of (non-)methylation and differences in the methylation level between groups can be determined by the differences in number of reads per locus per sample (Schield et al. 2016). The advantages of this technique resemble those of all genomic reduced representation approaches such as RADseq: the possibility of sampling genome wide, no requirement for a reference genome, and the ability to map loci against a reference genome (if available) to determine to which genomic region they correspond (Andrews et al. 2016; Schield et al. 2016).

Combining genetic and epigenetic analyses in the same study is to date underleveraged but particularly valuable for providing insight into the relationship between genetic and epigenetic variation and downstream effects of interest, such as phenotypic diversity (Hu & Barrett, 2017). For example, DNA methylation can explain phenotypic variation better than genetics (e.g. Richards, Schrey & Pigliucci, 2012), methylation pattern can be explained by genetic effects rather than by other variables of interest (Robertson et al. 2017), and population-level methylation analyses can provide insight to mechanisms of evolution (Gugger et al. 2016). To infer methylation and genomic polymorphism (SNPs) using separate NGS techniques for the same set of individuals is expensive, inefficient, and time consuming but is the approach that has been used to date (e.g. Dimond, Gamblewood, & Roberts, 2017). A combined molecular approach that allows for DNA methylation and genetic analyses would increase the efficiency of such approaches and increase the scope of possible research questions in this area and be of considerable value to this field of study.

Because epiRADseq is similar in molecular methodology to ddRADseq, in this study we test whether the SNPs recovered by epiRADseq can also be used for population genomics. If SNPs for population genetics can be reliably extracted from epiRADseq data then epigenetic and population genomic analyses can be conducted efficiently on the same samples using the same molecular technique, from DNA extraction through to library preparation and sequencing. We tested this using two independent examples from natural animal populations for which epiRADseq and ddRADseq data are available from the same individuals: a previously published dataset (Dimond et al. 2017) from a marine invertebrate, the corals of the genus *Porites* (genome size between 420 Mb and 1.14 Gb) for which there is currently no reference genome; and a newly generated dataset from a vertebrate, the freshwater European whitefish *Coregonus lavaretus* (genome size 3.3 Gb) for which genome scaffolds are available. We ran analyses in parallel on epiRADseq and ddRADseq data to compare number of SNPs retained, summary statistics, and inferred population genetic structure. We conclude that epiRADseq data are appropriate for population genomics and suggest a bioinformatic pipeline for extracting SNPs.

## Material and Methods

### 2.1 Coral data source

EpiRADseq was recently used in conjunction with ddRADseq by Dimond et al. (2017) to assess the population genetics and epigenetics of three morphospecies of coral *Porites spp*. from the Caribbean. EpiRADseq was used for differential methylation analysis and ddRADseq was used to estimate population structure between samples and to correct for the bias of epiRADseq in the methylation analysis, as a missing locus could either mean a lack of site due to mutation (a genetic factor) or due to methylation (an epigenetic factor) (Schield et al. 2016). They excluded from the dataset all epiRADseq loci that were missing in the ddRADseq dataset. However, they did not test the possibility of using epiRADseq to call SNPs for genetic analysis.

### 2.2 Coral data processing

The raw reads for ddRADseq and epiRADseq from Dimond et al. (2017) were downloaded from their repository (http://owl.fish.washington.edu/nightingales/Porites_spp/). The coral data comprised 48 individuals prepared with both ddRADseq and epiRADseq methods, for a total of 96 samples split into 12 libraries, of which we focused on the 60 samples (30 ddRADseq and 30 epiRADseq) that were analysed in the Dimond et al. (2017) study.

The first 5 and 3 bp were trimmed with *Trimmomatic* from the forward and reverse reads to remove the enzyme cut site. Then, paired-end trimming was done with following settings: LEADING = 20, TRAILING = 20, MINLEN = 85. Reads were mapped against the genome of the coral symbiont *Symbiodinium minutum*, provided in the Supplementary information of Dimond et al. (2017), to remove symbiont reads from the *de novo* assembly, as was done by Dimond et al. (2017), using *bwa mem* (Li & Durbin, 2009). The retained coral reads were used for all further analyses.

A pseudo-reference genome of coral samples was created so that we could determine the number of common SNPs found in both the ddRADseq and epiRADseq datasets. This pseudo-genome was assembled using *Rainbow* v.2.0.4 (Chong et al. 2012) with the cluster, divide, and merge functions with default parameters using the fastq files free of symbiont reads. *CD-Hit* v.4.7 (Fu et al. 2012) was then used with the cd-hit-est (at a 90% identity threshold) function for further filtering. ddRADseq and epiRADseq reads were mapped against this pseudo-genome using *bwa mem* with default settings and retained if mapping quality was >20.

If a sample had fewer than 200,000 reads in either the ddRADseq or epiRADseq dataset, it was removed from both so that the datasets had the same individuals. This excluded four samples and thus 52 samples (26 for ddRADseq and 26 for epiRADseq) were retained for analysis. The *ref_map.pl* v.2.1 pipeline in Stacks (Catchen et al. 2013) was run for both ddRADseq and epiRADseq using default parameters. All the samples were considered as part of the same population for the Stacks pipeline. The dataset was then filtered with the following parameters from the *populations* program: -*r* = 1 (no missing data allowed, same as in Dimond et al. (2017)), --*min_maf* = 0.10 and --*max_obs_het* = 0.6, --*write_single_snp*.

### 2.3 Whitefish data generation

Using existing tissue samples of *Coregonus lavaretus* from four Scottish loch populations preserved in ethanol (Crotti, Adams & Elmer, unpubl), DNA was extracted from fish fin clips for the ddRADseq and muscle tissue for the epiRADseq libraries using the NucleoSpin Tissue kit (Macherey-Nagel) following the manufacturers recommendations. The protocol used for the ddRADseq library preparation follows Jacobs et al. (2018). Briefly, 1 μg of genomic DNA per sample was double digested using the rare cutting enzyme *PstI*-HF (CATCAG recognition site) and the common cutting enzyme *MspI* (CCGG recognition site). Combinatorial barcoded Illumina adapters were then ligated to *PstI*-HF and *MspI* overhangs. Samples were size selected using a Pippin Prep (Sage Science) at a target range of 150-300 bp fragments. To enrich for the selected loci, we performed PCR amplification cycles with the following settings: 30 s at 98 °C, 9X (10 s 98 °C, 30 s 65 °C, 30 s 72 °C), 5 min 72 °C. After PCR purification, the library was run on a 1.25% agarose gel stained with SYBR Safe (Life Technologies) to remove any adapter dimers and/or fragments outside the selected size range. DNA was excised manually, cleaned and quantified using the Qubit Fluorometer with the dsDNA BR Assay (Life Technologies) to ensure the final library concentration of >1 ng/μL.

The protocol for the epiRADseq library was identical to the ddRADseq, except a methylation sensitive *HpaII* (CCGG recognition site; therefore, compatible with the same combinatorial barcodes and adapters) was used instead of the *MspI* restriction enzyme.

The ddRADseq and epiRADseq libraries consisted of the same 43 samples each, including two technical replicates to estimate sequencing error (Mastretta-Yanes et al. 2015), and were sequenced on a single lane to 4 million reads per individual. NGS sequencing was carried out at Glasgow Polyomics facility on the Illumina NextSeq 500 platform with 75 bp paired end reads.

### 2.4 Whitefish data processing

EpiRADseq and ddRADseq data were analysed separately using the same approaches. Samples with fewer than 350 K reads in one dataset were excluded from both datasets. The filtering steps applied to the whitefish data were similar as used in the coral data, but with some modifications because the whitefish data were analysed as single end. First, raw reads were demultiplexed with *process_radtags* in Stacks v.2.1 (Catchen et al. 2013) and only forward reads were retained. *Trimmomatic* (Bolger et al. 2014) was used to trim reads with following settings: HEADCROP = 5 (to remove enzyme cutting site), LEADING = 20, TRAILING = 20, MINLEN = 60. Reads were then mapped to an unpublished draft genome of the lake whitefish *Coregonus clupeaformis* (L. Bernatchez, pers comm) using *bwa mem* v.0.7.17 with default settings and retained if mapping quality was > 20 with *samtools* v. 1.7 (Li et al. 2009). In Stacks, the *ref_map.pl* script was used to assemble reads into stacks and call loci, and the *population* module was used to call SNPs.

To assess the sensitivity of SNP calling to missing data for epiRADseq data, we created three different datasets for both the ddRADseq and epiRADseq reads, which varied according to the proportion of individuals per population the locus had to be in to be retained (-*r* parameter): 0.667, 0.75, or 1. The other filtering parameters were kept constant: -*p* = 2, --*max_obs_het* = 0.6, --*min_maf* = 0.10, --*write_single_snp*. The three datasets are hereafter referred to as the -*r* 67, -*r* 75, and -*r* 100 datasets. This assessment was done only for the whitefish data, as with the coral data we focused on comparing our results to the original paper (Dimond et al. 2017).

We tested the effect of allele dropout (ADO) on genetic estimates derived from epiRADseq data, because methylated loci are not sequenced (Schield et al. 2016). To do so we estimated genetic diversity for each individual for both ddRADseq and epiRADseq data in *Stacks* with following parameters: -*p* 23 (each individual was considered a population), --*max_obs_het* = 0.6, --*min_maf* = 0.10. We then compared these estimates using a paired Wilcoxon signed rank test.

### 2.5 Whitefish and coral data analysis

For both the whitefish and coral data we recorded the total number of SNPs retained by ddRADseq and epiRADseq datasets. Summary statistics of genetic diversity (expected heterozygosity, observed heterozygosity, and nucleotide diversity) per locus calculated by the *population* module of Stacks for the ddRADseq and epiRADseq datasets were compared using Spearman correlation in the R environment (R Core Team, 2018).

To compare estimates of population structure between the ddRADseq and epiRADseq datasets, we used the R package *adegenet* v.2.1.1 (Jombart, 2008) to run a Discriminant Analysis of Principal Components (DAPC) (Jombart et al. 2010), which uses *k*-means clustering and the Bayesian information criterion to identify the most likely number of genetic clusters in the dataset. The *xvalDAPC* function was used to determine the number of PCs to be retained by the DAPC analysis. The divergence estimate between the inferred clusters was calculated using Weir and Cockerham Fst (Weir & Cockerham, 1984) implemented in the R package *hierfstat* v.0.04 (Goudet, 2005). For the coral analysis we additionally ran the DAPC on the set of SNPs used by Dimond et al. (2017), which they made available in the supplementary information of their article, to compare our results to the original study.

### 2.6 Genotyping error rate

To estimate genotyping error rate for the whitefish data we used two approaches: 1) we computed a matrix of genetic distances between individuals using the function *dist.gene* in the R package *ape* v.5.2 (Paradis & Schielp, 2018), following Dimond et al. (2017); 2) we used the R script published by Mastretta-Yanes et al. (2015), where the number of SNP mismatches is counted and calculated as the ratio over all compared loci (Recknagel et al. 2015). Replicated samples were compared at six-fold coverage. Technical replicates were not included in the coral dataset so genotyping error was not quantified.

## Results

### 3.1 Coral data filtering

The 30 ddRADseq samples had a total of 213 M raw reads and the 30 epiRADseq samples a total of 156 M raw reads (Table 1). After filtering with *Trimmomatic*, the ddRADseq samples retained 205 M reads, and the epiRADseq samples retained 149 M reads. Mapping against the pseudo-genome created from the ddRADseqs reads (418,401 contigs), retained 142 M reads for the ddRADseq and 102 M reads for the epiRADseq samples.

**Table 1.**
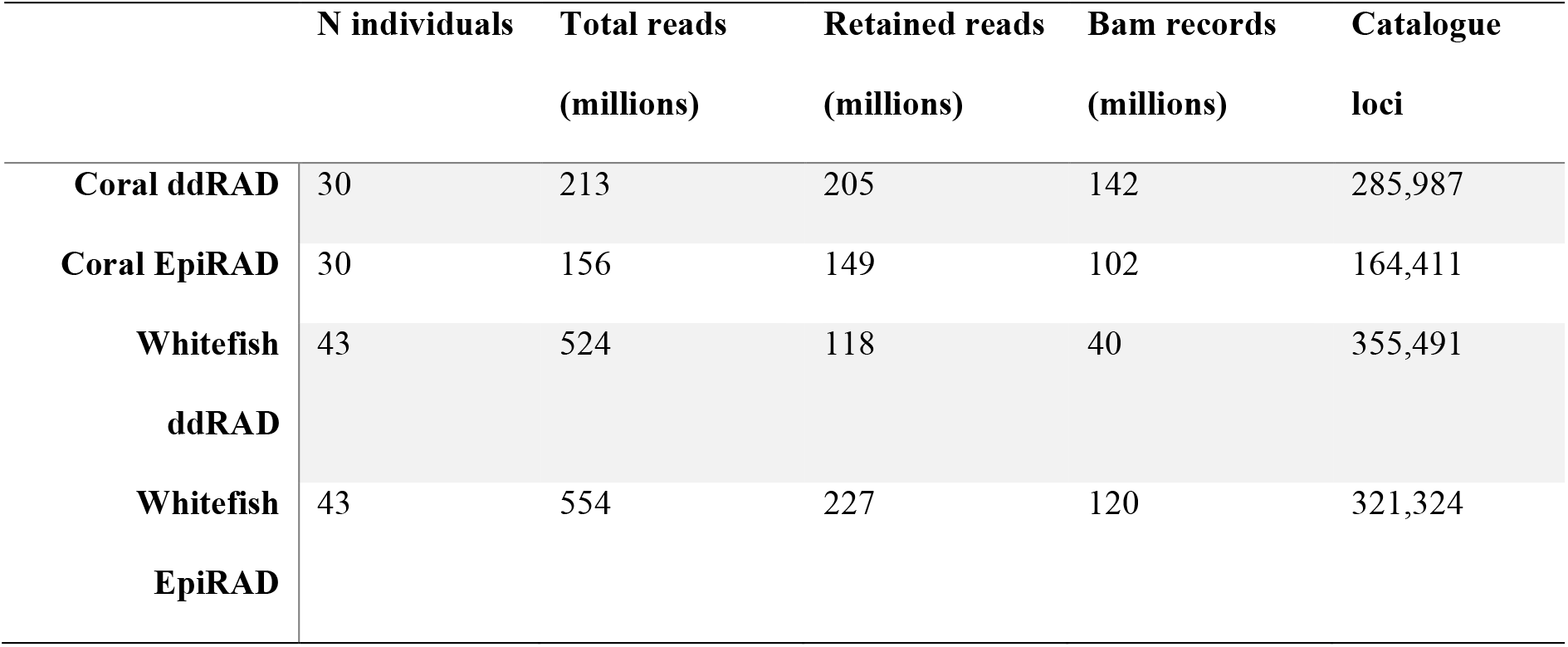
Number of samples in the libraries, and number of reads retained (in millions, M) after each step. Retained reads is the number after demultiplexing and Trimmomatic. BAM records refers to the number of reads retained after mapping to (pseudo)reference draft genome. Catalogue loci are the total loci inferred from Stacks, whether variable or not.

The *Stacks* pipeline generated a catalogue of 285,987 loci for the ddRADseq dataset, with a mean effective per sample coverage of 64.9x, and 164,411 loci for the epiRADseq dataset, with an effective per sample mean coverage of 75.7x. The average number of loci per individual was 58,896 for the ddRADseq and 33,843 for the epiRADseq catalogues.

### 3.2 Coral data analyses

The *population* filtering generated datasets of 1,046 SNPs and 819 SNPs for ddRADseq and epiRADseq respectively (Fig. 1a). The number of SNPs retained in our study is slightly lower to those used by the original study (1,113 SNPs from ddRADseq, also assessed here). By mapping reads to a reference assembly, we could calculate the number of SNPs that overlapped between the two datasets. In total 676 SNPs overlapped, which corresponds to 83% of SNPs in the epiRADseq and 65% of SNPs in the ddRADseq datasets.

**Figure 1.**
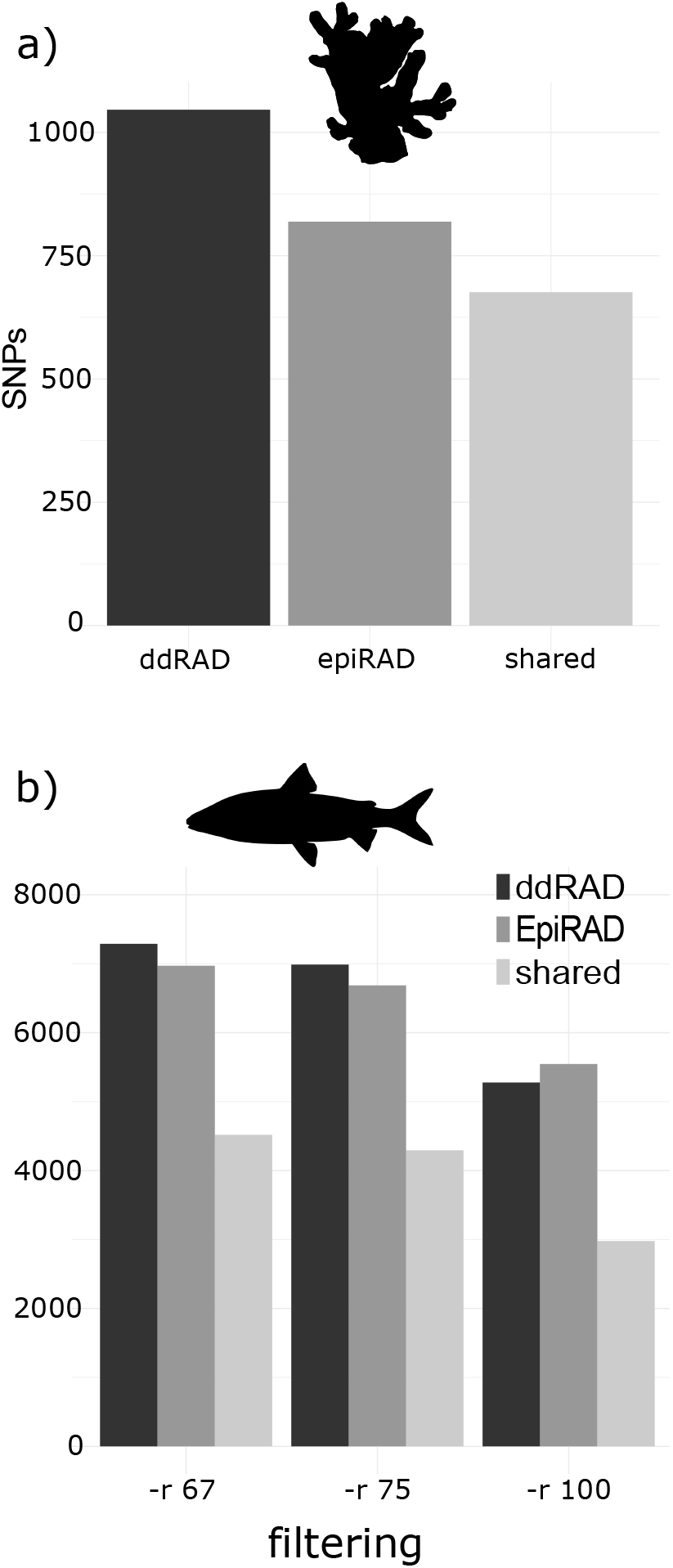
The number of SNPs retained by the ddRADseq and epiRADseq datasets for a) the coral data and b) whitefish data. Three datasets were created for the whitefish data, differing in the percentage of individuals that must possess a particular locus for it to be included (-*r* parameter of the *population* program from the *Stacks* pipeline).

DAPC analyses of the epiRADseq and ddRADseq datasets recovered the same three clusters as were inferred from the original study from Dimond et al. using ddRADseq (Fig. 2). Our Fst estimates between clusters ranged from 0.24 to 0.26, while the estimates of Dimond et al. were 0.19 to 0.21 (Fig 2a,b,c). The proportion of variation explained by the discriminant functions was similar in all three datasets (Fig. 2). When comparing estimates of genetic diversity, we recovered strong Spearman’s σ correlation for all three summary statistics between the ddRADseq and the epiRADseq datasets (Fig. 3).

**Figure 2.**
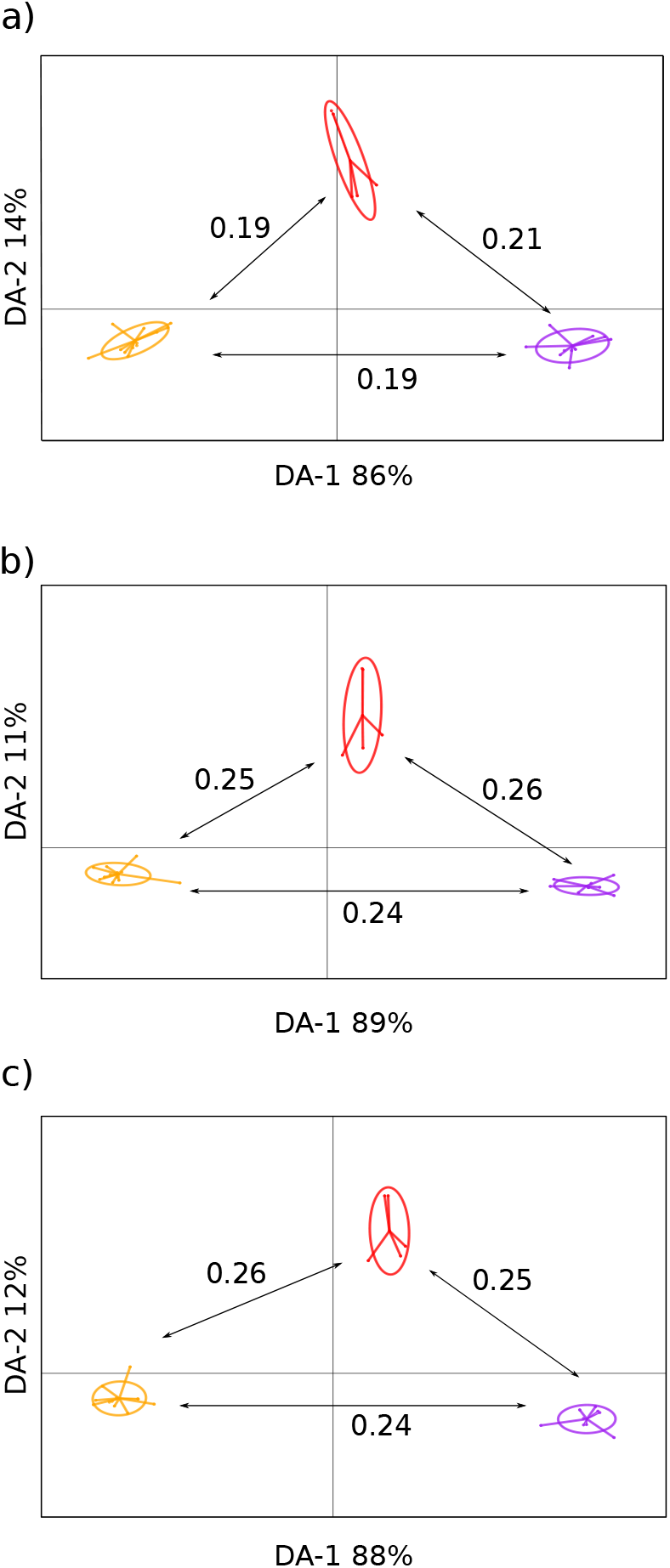
Results of the coral DAPC analyses of the a) SNPs used by Dimond et al. (2017), b) SNPs from the re-called ddRADseq dataset, and c) SNPs from the epiRADseq dataset. The analysis was based on five retained principal components, as suggested by the cross-validation of DAPC. These PCs were then summarised with two discriminant functions and percent variance captured appears on the axes. The numbers on arrows are Weir and Cockerham Fst values between the clusters.

**Figure 3.**
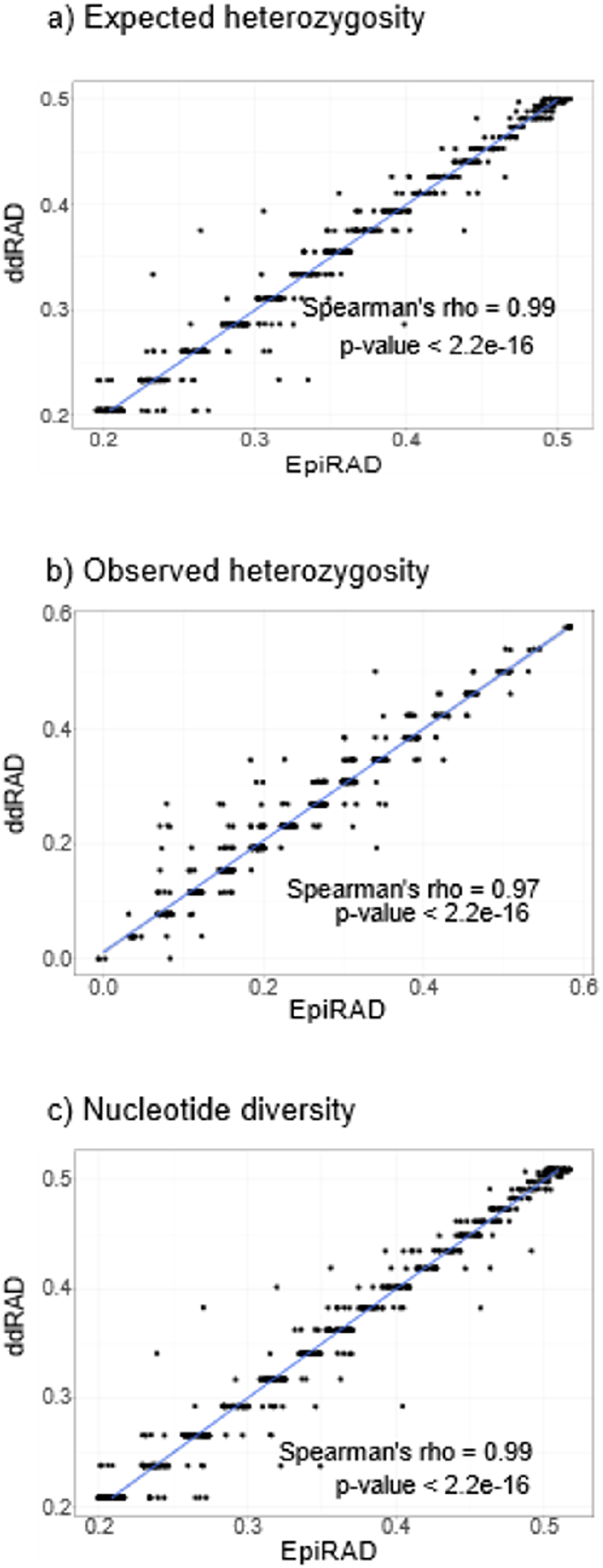
Correlation of a) expected heterozygosity, b) observed heterozygosity, and c) nucleotide diversity, between ddRADseq (y axis) and epiRADseq (x axis) estimates for the coral data. Each dot represents a genomic site from the “sumstats.tsv” file of the Stacks pipeline that was shared between the ddRADseq and the epiRADseq datasets.

### 3.3 Whitefish sequencing results and data filtering

The whitefish ddRADseq library generated a total of 524 M reads and the epiRADseq library generated 554 M reads (Table 1). After demultiplexing with *process_radtags* and filtering with *Trimmomatic*, the ddRADseq library retained 118 M reads, while the epiRADseq library retained 227 M reads. After mapping to the reference genome, the ddRADseq library retained 40 M reads, while the epiRADseq library retained 120 M reads (Table 1). Excluding the samples with fewer than 350 K reads left a total of 23 samples plus two technical replicates in the epiRADseqs dataset and 23 samples plus two technical replicates in the ddRADseq dataset.

The *Stacks* pipeline produced a catalogue of 355,491 loci for the ddRADseq library, with a mean effective per sample coverage of 12.7x, and of 321,324 loci for the epiRADseq library, with a mean effective per sample coverage of 36x. The average number of loci per individual was 108,127 and 110,614 for the ddRADseq and epiRADseq respectively.

### 3.4 Genotyping error rate

The SNP genotyping error rate in the whitefish dataset was lower for epiRADseq for both analysis approaches. The *dist.gene* approach recovered a mean error rate of 6% (± standard deviation 0.6%) for the ddRADseq, and of 3% (± 0.5%) for the epiRADseq, while the Mastretta-Yanes et al. approach estimated a mean error of 5% (± 0.3%) for the ddRADseq and of 3% (± 0.4%) for the epiRADseq.

### 3.5 Whitefish data analysis

The number of SNPs retained was very similar for those generated with the epiRADseq method and the ddRADseq method and decreased with increasing filtering stringency (Fig. 1); for the epiRADseq generated data we recovered 6971, 6686, and 5546 SNPs in the -*r* 67, -*r* 75, and -*r* 100 datasets respectively, while for the ddRADseq generated data we recovered 7289, 6988, and 5277 SNPs in the three datasets respectively. A total of 4518 SNPs were shared between the two -*r* 67 datasets, 4294 SNPs were shared between the two -*r* 75 datasets, and 2978 SNPs were shared between the -*r* 100 datasets.

The estimates of heterozygosity and nucleotide diversity inferred from ddRADseq and epiRADseq derived SNPs were highly correlated, with Spearman’s correlations of 88.5 to 92.8% (Table 2). When looking at the genetic diversity estimates per individual, which would be impacted by allele dropout, we observed no reduction in expected heterozygosity (V = 58, p-value = 0.45) or nucleotide diversity (V = 82, p-value = 0.31) for the epiRADseq data.

**Table 2.** Spearman’s correlation between coral and whitefish epiRADseq and ddRADseq estimates of expected heterozygosity (He), observed heterozygosity (Ho), and nucleotide diversity (Pi) for -*r* 67, -*r* 75, and -*r* 100 datasets. Number of sites corresponds to the SNPs shared between epiRADseq and ddRADseq datasets, for which the correlation was calculated.

|  | Stacks<br>filtering | Number of<br>sites | He | Ho | Pi |
| --- | --- | --- | --- | --- | --- |
| **Whitefish** | -r 67 | 4518 | 0.904 | 0.885 | 0.896 |
|  | -r 75 | 4294 | 0.911 | 0.889 | 0.903 |
|  | -r 100 | 2978 | 0.928 | 0.906 | 0.919 |
| **Coral** | -r 100 | 676 | 0.988 | 0.972 | 0.988 |

The results of the population genetic structure analysis with DAPC were consistent across filtering stringencies and datasets (Fig. 4), with the four populations being grouped into two genetic clusters separating on axis 1 (and so displayed on one axis of variation instead of the two shown for the corals). Fst divergence between the two clusters was identical between methods for the -*r* 67 and -*r* 75 datasets at Fst = 0.23, and it was negligibly higher for the ddRADseq in the -*r* 100 datasets at 0.24 and 0.25 (Fig. 4).

**Figure 4.**
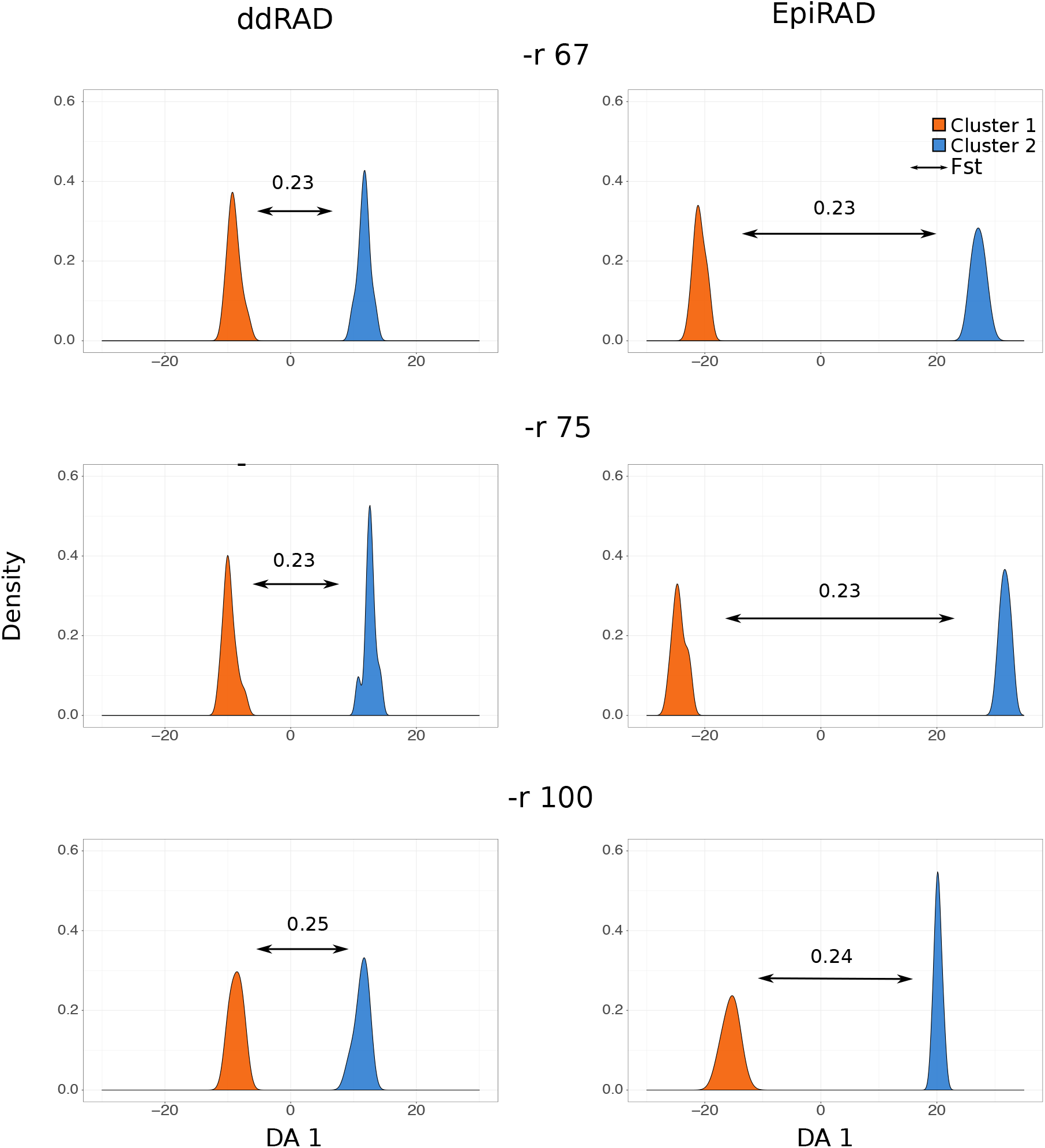
Results of the European whitefish DAPC analyses at three different filtering stringencies (-*r* 67, -*r* 75, -*r* 100). The analysis was based on five retained principal components, as suggested by the cross-validation of DAPC. These PCs were then summarised on one discriminant function, as only two genetic clusters are observed. The numbers above arrows represent Weir and Cockerham Fst values between the two identified clusters.

## Discussion

Here we used two independent natural animal population datasets to show that epiRADseq data can be used to derive SNPs for population genomic analyses. We compared SNP number, estimates of summary statistics, and inference of population structure between ddRADseq and epiRADseq methods in a newly generated dataset of European whitefish and a previously published dataset on corals. Overall, we find strong agreement for all of the above metrics between epiRADseq and ddRADseq protocols, meaning that epiRADseq data give equivalent results to the well-established method of ddRADseq-derived SNPs. The implication is that a single dataset can be used for epigenetic analyses and for inference of population structure. This is not only efficient but also valuable studies on the association between epigenetic and genetic diversity and their impact on phenotype.

Here we used previously published data and new data when comparing the epiRADseq and ddRADseq generated SNPs, which allows us to demonstrate the robustness of the molecular methods and of the bioinformatics pipelines independently. The coral dataset was drawn from Dimond et al. (2017), where they investigated population structure between three morphospecies of coral with ddRADseq and looked at the relationship between DNA methylation and environmental factors. The number of SNPs in our datasets is slightly lower than those used in the Dimond et al. (2017) study; we recovered 1,046 SNPs for ddRADseq and 819 SNPs for epiRADseq while they previous study retained 1,113 SNPs from ddRADseq. This is likely because different bioinformatic pipelines applied as they used *Pyrad* (Eaton, 2014) while we used *Stacks* (Catchen et al. 2013).

Our genetic diversity, differentiation, and population structure results of the coral data, derived from SNPs from their epiRADseq data, are consistent with those obtained by Dimond et al. (2017). The Fst estimates between the three population genetic clusters are slightly higher in our study (approximately 20% in excess of the previously published values). This is likely caused by the different loci being retained by the *Stacks* vs *Pyrad* pipelines, consistent with Pante et al. (2015) reporting a locus overlap of less than 50% between methods. However, Fst results are rarely strictly comparable across studies and instead are relative to the markers used (Hartl & Clark, 2007) and therefore these deviations can be considered irrelevant. These explorations and comparisons of our pipeline on the coral dataset demonstrate the appropriateness of the pipelines we applied and that the baseline genetic information is comparable across studies.

For the coral dataset, the number of loci in the ddRADseq catalogue was 43% higher than in the epiRADseq catalogue (285,987 vs 164,411) and resulted in a higher number of SNPs in the final ddRADseq dataset. This is expected due to the loci sampling bias of epiRADseq, as loci that are methylated are not sequenced (Schield et al. 2016). However, we show that there is negligible effect on the resulting summary and differentiation statistics and the epiRADseq SNPs are therefore equivalent to the ddRADseq SNPs.

We explored the effect of different filtering levels on the SNP retention of epiRADseq and ddRADseq derived SNPs from the whitefish data. We did not explore this with the coral data as we were more interested in comparing estimates of population structure between epiRADseq and the previously published estimates derived from ddRADseq. As expected, the -*r* 67 and -*r* 75 ddRADseq datasets had more SNPs than the respective epiRADseq datasets, but the epiRADseq -*r* 100 dataset had more SNPs than the ddRADseq -*r* 100 dataset. This is probably due to the higher coverage of the epiRADseq reads (85 M reads for 25 individuals in the epiRADseq vs 32 M reads for 25 individuals in the ddRADseq), which resulted in more SNPs being retained in the most stringently filtered dataset.

We find an agreement between ddRADseq and epiRADseq analyses of population structure in the whitefish data, as both methods recover two clusters in our dataset of four sampled and closely related populations. The -*r* filtering had some impact on the correlation of the summary statistics between ddRADseq and epiRADseq, with the correlation increasing from as low as 88% up to 92% as the filtering became more stringent. This is expected because of the -*r* parameter in *Stacks*, which influences the number of individuals in a population a locus must be present to be retained in the dataset. In the -*r* 67 and -*r* 75 datasets, it is not required for the locus to be present in the same set of individuals (i.e. in two-thirds or three-quarters of all individuals in a population, respectively), while in the -*r* 100 datasets this restriction is complete so all retained SNPs have to be shared across all individuals. We did not explore further filtering in our analyses, but previous work (e.g. Paris, Stevens, & Catchen, 2017; O’Leary et al. 2018; Linck & Battey, 2019) highlights the importance of fine-tuning the SNP-calling pipeline to suit the researcher’s needs and the specificity of each dataset. However, with regard to the use of SNPs from epiRADseq it is important to consider that comparability across different datasets is not what matters; here that is done to evidence the method. Instead each of these stringencies and datasets would be valid. Overall, these results suggest that allowing some missing data (i.e. −*r* of 67% or 75%) will not bias genetic analyses conducted with SNPs from epiRADseq data, consistent with what has already been shown previously with ddRADseq (Shafer et al. 2017).

We tested whether allele drop out (ADO) due to locus methylation (Schield et al., 2016) had an effect when using epiRADseq derived SNPs for genetic analyses. It has been shown through simulations (Gautier et al. 2013) and observed in empirical studies (Luca et al. 2011) that ADO leads to an underestimation of expected heterozygosity and nucleotide diversity. This could be a concern for epiRADseq derived SNPs because, by design, a methylated locus is not cut with epiRADseq and therefore will be absent from the dataset. However, we found no difference between ddRADseq and epiRADseq genetic diversity estimates per individual, suggesting ADO is not a particular concern in epiRADseq data.

Genotyping error in NGS techniques is due to several factors, including sequencing errors, assembly errors and missing data and will be influenced by coverage (Mastretta-Yanes et al. 2015). Using technical replicates is a way to estimate this error, which can then be moderated by fine-tuning the bioinformatic pipeline. We find that the SNP genotyping error rate is low and very similar between ddRADseq and epiRADseq libraries, ranging between 3 and 6% according to the calculation method used. Mastretta-Yanes et al. (2015) found SNP error rates between 2.4 and 5.8% using the *Stacks* pipeline on Illumina-based RAD sequencing. Recknagel et al. (2015), using a similar lab protocol to that used for the whitefish libraries here but sequenced on an Ion Proton platform, recovered genotyping errors of 1.8-2.2%. Dimond et al. (2017) used the ddRADseq and epiRADseq samples as technical replicates, as they were sequenced on the same lanes, and recovered a mean genotyping error rate of 3.6% (standard deviation 3.1%). Therefore, genotyping error rates in the whitefish libraries are consistent with those found by previous studies and are very similar between the ddRADseq and epiRADseq approaches.

When looking at the results of the coral and whitefish data together, we find agreement when estimating population structure either with ddRADseq or with epiRADseq. However, the percentage of SNPs shared between ddRADseq and epiRADseq was higher in the coral data (83% vs 55-65%). This could be due to the difference in genome complexity and genome size of the two organisms studied. Salmonids have undergone an extra whole genome duplication compared to other teleosts (Macqueen & Johnston, 2014) and members of the genus *Coregonus* have an estimated genome size of 3.3 Gb (Gregory, 2018). Members of the coral order Scleractinia, to which the coral genus *Porites spp*. belong, have genomes ranging from 420 Mb to 1.14 Gb (Gregory, 2018). Smaller genomes generate fewer RAD loci, which are then more likely to be found across sequencing libraries at a given coverage (see Recknagel et al. 2015 for detailed quantifications). Furthermore, DNA methylation levels and patterns differ between the organisms studied here and may have an impact. Most of the CpG sites (~80%) in vertebrate genomes are methylated, with the unmethylated sites forming regions known as CpG islands, which are usually located near gene promoters (Metzger & Schulte, 2016). In contrast, most of the methylation in invertebrates occurs specifically in CpG sites within gene bodies (Dixon, Bay, & Matz, 2014). The methylation level of CpG sites in the scleractinian coral *Stylophora pistillata* is around 7% (Liew et al. 2018), a stark contrast to the methylation level of vertebrates. Differences in methylation between organisms might influence the number of fragments that are cut during digestion with *HpaII* and therefore affect the number of loci sequenced. We did not explore the genomic location of the SNPs used here, but with appropriate reference genome annotation information that is possible and would be very informative.

In addition to EpiRADseq (Schield et al. 2015), other methylation-sensitive techniques have been developed to take advantage of the basic RADseq methodologies. MethylRAD (Wang et al. 2015) is based on the 2b-RAD methodology (Wang et al. 2012) and employs methylation sensitive Mrr-like enzymes that, like IIB restriction enzymes, cut upstream and downstream of the recognition site if it is methylated. Instead, enzymes used for ddRADseq and epiRADseq only cut downstream of the recognition site. Like epiRADseq, this technique does not provide base-pair resolution of methylation but provides methylation information by comparing locus read depth across samples to infer abundance. Given its similarity to 2b-RAD, we suspect that MethylRAD could also be used for extracting SNPs for genetic analyses as well, although thorough testing should be carried out. BsRADseq (Trucchi et al. 2016) combines RADseq with bisulfite sequencing, providing a base pair-resolution of DNA methylation, similarly to RRBS. We also believe this technique could be used for both genetic and epigenetic analyses, but again we recommend testing to explicitly compare the genotype datasets.

Here, we showed that the recently developed epiRADseq approach for the study of DNA methylation variation can also be used for generating SNPs for population genetic analyses, using both reference-based and *de novo* approaches. Sequencing only an epiRADseq library halves the cost in time, consumables, and sequencing compared to sequencing ddRADseq for SNPs and epiRADseq for methylation abundance. This combination provides informative biological data for population genomics and differential methylation, which is a topic of growing interest in molecular ecology and evolution for its heritable and non-heritable effects (Hu & Barrett, 2017).

## Acknowledgements

This work was funded by Univ. of Glasgow College of Medical, Veterinary and Life Sciences doctoral training programme. We thank JL Dimond, SK Gamblewood, and SB Roberts for making their data public so we could explore it for this paper. We thank Glasgow Polyomics and J Galbraith for sequencing, M Capstick for support in the laboratory, and A Jacobs for analysis advice and comments on the draft manuscript. For access to unpublished lake whitefish scaffolds, we thank L Bernatchez, C Rougeux, S Pavey, E Normandeau, S Lien, and T Nome. We declare no conflict of interest.

## Data Accessibility

Data will be archived and made available in University of Glasgow Enlighten Repository with manuscript acceptance.

